# Molecular Evolution of Podocyte Slit-diaphragm Proteins

**DOI:** 10.1101/2020.11.03.366161

**Authors:** NSK Mulukala, V Kambhampati, SAH Qadri, AK Pasupulati

## Abstract

Vertebrates kidneys contribute to the homeostasis by regulating electrolyte, acid-base balance, and prevent protein loss into the urine. Glomerular podocytes constitute blood-urine barrier and podocyte slit-diaphragm, a modified tight junction contributes to the glomerular permselectivity. Nephrin, podocin, CD2AP, and TRPC6 are considered to be crucial members, which largely interact with each other and contribute to the structural and functional integrity of the slit-diaphragm. In this study, we analyzed the distribution of these four-key slit-diaphragm proteins across the organisms for which the genome sequence is available. We found that nephrin has a diverse distribution ranging from nematodes to higher vertebrates whereas podocin, CD2AP, and TRPC6 are predominantly restricted to the vertebrates. In the invertebrates nephrin and its orthologs consist of more immunoglobulin-3 and immunoglobulin-5 domains when compared to the vertebrates wherein, CD80-like C2-set Ig2 domains were predominant. Src Homology-3 (SH3) domain of CD2AP and SPFH domain of podocin are highly conserved among vertebrates. Although the majority of the TRPC6 and its orthologs had conserved ankyrin repeats, TRP, and ion transport domains, the orthologs of TRPC6 present in *Rhincodon typus* and *Acanthaster planci* do not possess the ankyrin repeats. Intrinsically unstructured regions (IURs), which are considered to contribute to the interactions among these proteins are largely conserved among orthologs of these proteins, suggesting the importance of IURs in the protein complexes that constitute slit-diaphragm. This study for the first time reports the evolutionary insights of vertebrate slit-diaphragm proteins and its invertebrate orthologs.

## Introduction

The vertebrate kidneys regulate the homeostasis of the body by maintaining fluid and acid-base balance, in addition to getting rid of toxic metabolic byproducts [1]. Nephron, the functional unit of the vertebrate kidney consists of glomerulus and tubule, wherein the former ensures ultrafiltration of plasma and the later regulates selective absorption of the glomerular filtrate [1]. These two units of nephron work in concert and regulates the final composition of urine. The three anatomical layers that constitute glomerular filtration apparatus are glomerular endothelium, glomerular basement membrane, and podocytes. Podocytes are visceral epithelial cells of the glomerulus and they seek greater attention owing to their unique localization and their crucial role in the glomerular biology. Podocytes are highly specialized cells with unique morphology and large nucleus to cytoplasmic ratio. Primary processes of podocytes ultimately branch into regularly spaced foot-processes that enwrap and provide epithelial coverage to the blood capillaries. Interdigitating podocyte foot-processes form modified junction called the slit-diaphragm (SD) [2]. SD serves as a size and shape-selective barrier, preventing plasma proteins to filter into urine thus curb the protein loss. Proteins such as nephrin, podocin, CD2AP, and TRPC6 are major components of the SD and contribute to the integrity of SD and ensure the ultrafiltration of urine.

In podocytes, the SD develops initially as a tight junction during the formation of comma and S-shape stages of the glomerular development [3]. Eventually as the glomerular development progresses, the primary SD structure evolves into a modified tight-adherens junction [4]. It is observed that proteins from both tight and adherens junction co-localize at the SD alongside neuronal junction proteins such as nephrin, Kin of IRRE like-1 (KIRREL1), etc [4]. The SD width ranges from 20-50nm which is sufficient to curb the passage of proteins from blood into urinary space [5]. Apart from helping in primary filtration, the SD also acts as a complex signaling hub. Preliminary evidence suggests that proteins: nephrin, KIRREL1, podocin, and TRPC6 participate in the signaling events that dictate the podocyte morphology [2].

Altered podocyte morphology (effacement) leads to excess proteinuria, hypoalbuminemia, and edema which are hallmark symptoms of nephrotic syndrome (NS) [6]. Corticoid therapy is the usual recourse to abate NS. However, patients that do not respond to the corticoid therapy suffer from steroid resistant form of NS (SRNS). Patients with congenital podocytopathy usually fall into SRNS category and display mutations in the proteins that constitute the SD. Nephrin encoded by *NPHS1* is a critical structural component of the SD and bridges the distance between interdigitating foot-processes of the podocytes. Nephrin was identified as a product of the mutated gene in patients with Finnish-type nephrotic syndrome. Nephrin has a long extracellular domain containing immunoglobulin (Ig)-like modules and a cytoplasmic fibronectin type III (FN3) domain [5]. Podocin is a stomatin family membrane protein and an important component of the SD complex. Stomatin family consists of evolutionarily conserved of Stomatin, Prohibitin, Follitin, and HflC (SPFH) aka Prohibitin (PHB) domain found from bacteria to mammals [7]. Podocin is encoded by *NPHS2*, which is frequently mutated and contributes to numerous SRNS cases [8]. CD2-associated protein (CD2AP) interacts with nephrin and podocin and localizes to the cytoplasmic face of the SD [9]. Transient receptor potential channel-6 (TRPC6) is another significant molecule that localizes to the SD. TRPC6 belongs to the larger family of TRP proteins [10]. TRPC6 interacts both with nephrin and podocin, suggesting that the regulation of calcium signaling mediated by TRPC6 is highly associated with the maintenance of the SD function. Although these four proteins are extensively discussed and form the crux of SD architecture several other proteins such as KIRREL1, P-cadherin, and FAT1 are also localized to the SD of podocytes.

Although invertebrates do not possess typical nephrons, nephron-like components can be found in the excretory systems of many invertebrates, indicating that the complexity of the vertebrate excretory system was inherited from their invertebrate systems. For example, the insect nephrocytes and the nephrons in the human kidney share several similarities [11, 12]. Furthermore, in *Drosophila melanogaster*, the orthologues of the major constituents of the SD were expressed in the nephrocytes and form a complex that closely mirrors the vertebrate SD complex [12]. The similarities between invertebrate nephrocytes and vertebrate podocyte suggest that these cell types are evolutionarily related. Relations such as this tingles the interest to investigate the evolution of the SD proteins and find the relevant orthologs in different metazoans. Our study is aimed to identify the orthologs of the nephrin, CD2AP, podocin, and TRPC6 across metazoans. We analyzed the domain composition and intrinsically unstructured regions (IURs) of the identified proteins to assess the relationship and the evolution of these orthologous proteins with the human SD proteins.

## Materials and Methods

### Identifying the orthologues

Four human SD proteins namely nephrin (NCBI: NP_004637.1), CD2AP (NCBI: NP_036252.1), podocin (NCBI: NP_055440.1), and TRPC6 (NCBI: NP_004612.2) were used to identify the orthologs in metazoan organisms with complete genome sequence. The reciprocal best hit approach with default settings in the pBLAST tool of the NCBI database was used to identify the orthologs. In cases where an ortholog of a protein could not be identified, additional searches were performed to identify potential orthologs against the next closest organism or phylum. The analysis was performed with the annotated protein sequences from 27 completely sequenced genomes.

### Protein alignment and phylogenetic analysis

Multiple sequence alignment (MSA) of the retrieved orthologous proteins FASTA sequences from the NCBI database was performed using Multiple Sequence Comparison by Log-Expectation (MUSCLE) tool of the MEGA X software. Default parameters such as gap penalty (2.90), hydrophobicity multiplier (1.20), and UPGMA cluster method were used to perform MSA. Phylogenetic trees were also derived for nephrin, CD2AP, podocin, and TRPC6 using the results from the respective MSAs. The maximum-likelihood method was used for constructing the phylogenetic trees using the default parameters in the MEGA-X software.

### Domain Analysis

Various domains in the identified orthologs were predicted using the protein families (Pfam) database with an E-value cut-off of 1.0. Visual representation of the domain in the sequences was done using the illustrator of biological sequences (IBS) software ver. 1.0. Further, we analyzed the domain sequences using the PSI-PRED server which uses a two-stage neural network to predict the secondary structure using data generated by BLAST. Furthermore, the server also extrapolated intrinsically unstructured regions (IUR’s). According to the PSI-PRED server a prediction confidence ≥0.5 indicates the residues as intrinsically unstructured. The IURs in the organisms were plotted as scattered plots using the OriginPro 2020 software.

## Results

### SD proteins are confined to vertebrates with fewer exceptions

We used the reciprocal best hit method to find the orthologs for human SD proteins in both the invertebrate and vertebrate phyla. The list of human SD proteins and their orthologs in various invertebrate and vertebrate phyla is tabulated along with the respective sequence accession numbers (Table.1). Our analysis revealed that nephrin is present in several organism ranging from Tardigrada to higher vertebrates suggesting a diverse distribution (Fig.1A). In the case of nematodes and arthropods we observed synaptogenesis protein-2 (*Caenorhabditis)* and sticks and stones protein (*Drosophila)* as the orthologs for nephrin. However, we could not identify the nephrin orthologs in the phyla Onychophora, Nemertea, Phoronida, Cyclostomata, and Chondrichthyes. It is interesting to note that Aves despite possessing the SD structure do not have nephrin or any orthologs of nephrin. We observed nephrin orthologs in the phyla Platyhelminths, Annelida, and Cephalochordata. Nevertheless, due to the limited genome sequence data availability we could not identify the nephrin ortholog names in the above-mentioned phyla. Unlike nephrin; CD2AP, podocin, and TRPC6 were majorly restricted to the vertebrate phylum with few exceptions. We noticed that only CINDR (CIN85 and CD2AP related) protein in arthropods showed homology with CD2AP (Fig.1B). Similarly, the TRPC3 protein in Mollusca and Echinodermata and TRPC7-like protein in Hemichordates were found to be closely related to TRPC6 (Fig.1C). Although TRPC6 was predominantly restricted to vertebrates, it is also identified in Rotifers. Our analysis revealed that podocin was present only the vertebrate organisms and we did not find podocin or its related proteins outside the vertebrate phylum (Fig.1D). These results suggest that although the distribution of nephrin is diverse, CD2AP, podocin, and TRPC6 are majorly restricted to vertebrates.

**Table 1:** Human slit-diaphragm proteins and its orthologs proteins NCBI accession IDs identified by reciprocal best hit method. *Note*: ‘§’indicates the ortholog of the respective SD protein identified from the organism mentioned in the braces.

| Phylum | Organism | Tax ID | Nephrin | CD2AP | Podocin | TRPC6 |
| --- | --- | --- | --- | --- | --- | --- |
| Priapulida | *Priapulus caudatus* | 37621 | XP_01466708<br>8.1 | No relevance | No relevance | No relevance |
| Nematode | *Caenorhabditis elegans* | 6239 | NP_00130967<br>4.1 | No relevance | No relevance | No relevance |
| Tardigrada | *Hypsibius dujardini* | 232323 | OQV25698.1 | No relevance | No relevance | No relevance |
| Onychophora | *Euperipatoides rowelli* | 49087 | No relevance | No relevance | No relevance | No relevance |
| Arthropoda | *Drosophila melanogaster* | 7227 | NP_788286.1 | NP_00126312<br>9.1 | No relevance | No relevance |
| Rotifera | *Brachionus plicatilis* | 10195 | RNA22476.1 | No relevance | No relevance | RNA42410.1<sup>§</sup><br>(*Xenopus tropicalis*) |
| Platyhelminthes | *Macrostomum lignano* | 282301 | PAA62947.1 | No relevance | No relevance | No relevance |
| Mollusca | *Octopus bimaculoides* | 37653 | XP_01477668<br>8.1 | No relevance | No relevance | XP_01478321<br>8.1<sup>§</sup><br>(*Xenopus tropicalis*) |
| Annelida | *Capitella teleta* | 283909 | ELU18150.1 | No relevance | No relevance | No relevance |
| Nemertea | *Notospermus geniculatus* | 416868 | No relevance | No relevance | No relevance | No relevance |
| Brachiopoda | *Lingula anatina* | 7574 | XP_01338021<br>3.1 | No relevance | No relevance | No relevance |
| Phoronida | *Phoronis australis* | 115415 | No relevance | No relevance | No relevance | No relevance |
| Hemichordate | *Saccoglossus kowalevskii* | 10224 | XP_00681964<br>7.1 | No relevance | No relevance | XP_00273056<br>9.1<sup>§</sup><br>(*Danio rerio*) |
| Echinodermata | *Acanthaster planci* | 133434 | XP_02211042<br>3.1 | No relevance | No relevance | XP_02210360<br>6.1 |
| Cephalochordata | *Branchiostoma floridae* | 7739 | XP_00259012<br>1.1 | No relevance | No relevance | XP_00260743<br>4.1<sup>§</sup><br>(*Xenopus tropicalis*) |
| Urochordata | *Ciona intestinalis* | 7719 | XP_00212274<br>7.1 | No relevance | No relevance | No relevance |
| Cyclostomata | *Petromyzon marinus* | 7757 | No relevance | No relevance | No relevance | No relevance |
| Chondrichthyes | *Rhincodon typus* | 259920 | No relevance | XP_02037068<br>5.1 | XP_02038250<br>9.1 | XP_02038393<br>8.1 |
| Osteichthyes | *Danio rerio* | 7955 | XP_01720650<br>3.1 | NP_00100858<br>3.2 | NP_00101815<br>5.2 | XP_00516124<br>7.1 |
| Amphibia | *Xenopus tropicalis* | 8364 | XP_03176155<br>2.1 | NP_00112143<br>5.1 | XP_01794909<br>6.1 | XP_00293561<br>6.2 |
| Reptilia | *Anolis* | 28377 | XP_01685151 | XP_00811479 | XP_00322550 | XP_00810627 |
|  | *carolinensis* |  | 4.1 | 6.1 | 6.1 | 3.1 |
| **Aves** | *Gallus gallus* | 9031 | **Absent** | NP_00130533<br>2.1 | XP_422265.3 | XP_417184.4 |
| **Mammalia** | *Ornithorhynchus anatinus* | 9258 | XP_02892103<br>4.1 | XP_02892802<br>7.1 | XP_00151573<br>4.2 | XP_02890356<br>5.1 |
|  | *Rattus norvegicus* | 10116 | NP_072150.1 | AAM47029.1 | NP_570841.2 | NP_446011.1 |
|  | *Mus musculus* | 10090 | NP_062332.2 | AAI38375.1 | NP_569723.1 | XP_00650991<br>2.1 |
|  | *Pan troglodytes* | 9598 | XP_01679121<br>6.2 | XP_00944969<br>0.1 | XP_01678887<br>5.1 | XP_01677734<br>1.2 |
|  | *Homo sapiens* | 9606 | NP_004637.1<br>(Gene ID:<br>4868) | NP_036252.1<br>(Gene ID:<br>23607) | NP_055440.1<br>(Gene ID:<br>7827) | NP_004612.2<br>(Gene ID:<br>23607) |

**Figure 1:**
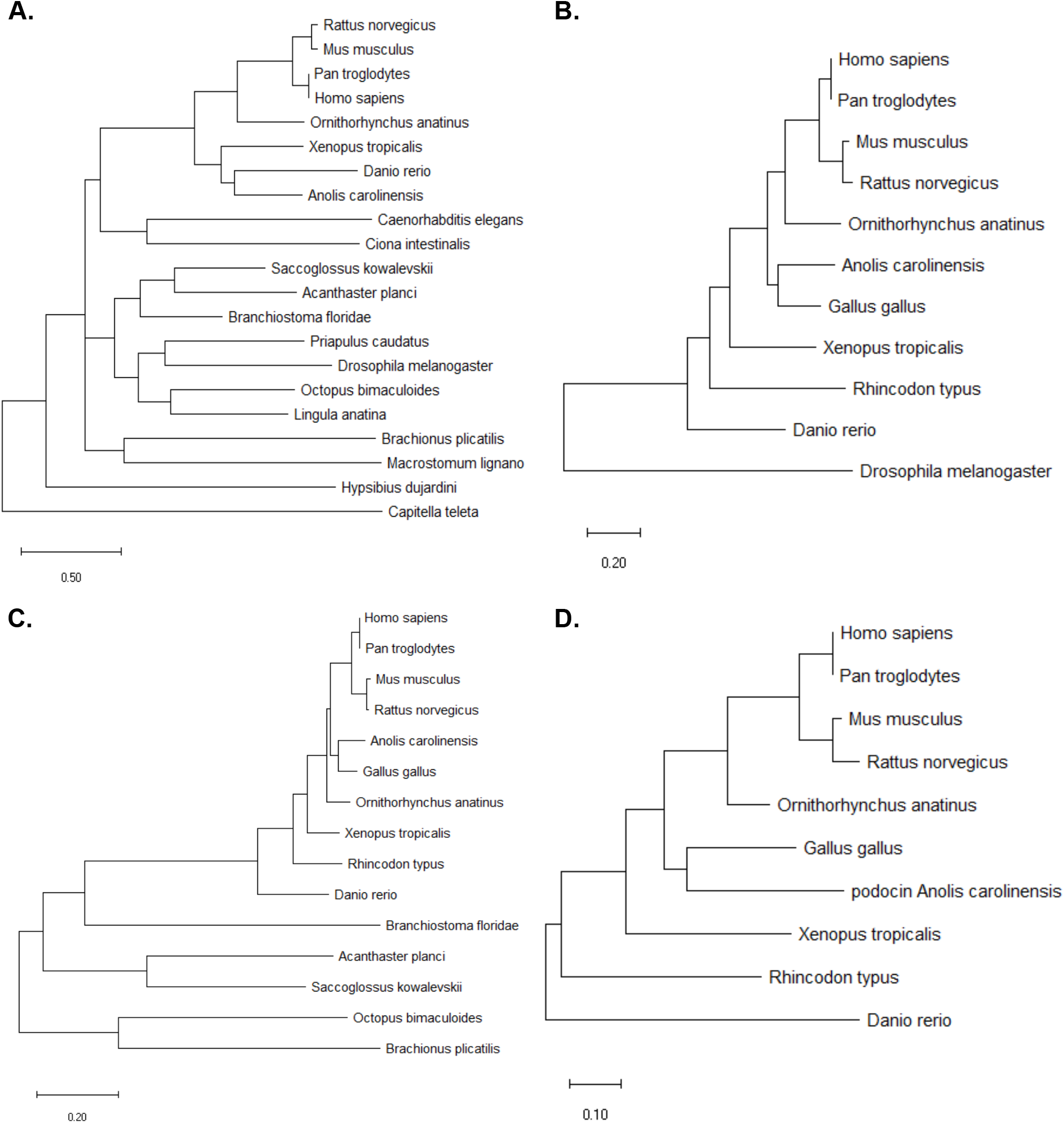
Distribution of key slit-diaphragm proteins and its orthologs across the vertebrates and invertebrates: A) nephrin, B) CD2AP, C) podocin, and D) TRPC6. Organisms with complete genome sequence were considered for the study.

### SD proteins and its orthologs share conserved domains

As we observed the presence of the SD proteins and their orthologs in various phyla predominantly from vertebrates, we next assessed the evolutionary accumulation and conservation of unique domains in these proteins. Human nephrin consists of eight immunoglobulin (Ig) domains and a fibronectin-3 (FN-3) domain [13]. Based on the sequence and number of strands in the β-sandwich of the Greek key motif (of Ig domain) we observed that human nephrin consists of one Ig5, five-CD80-like C2-set Ig2, two-Ig3 domains (Fig.2). Our analysis revealed that in the invertebrates both nephrin and its orthologs consist of more Ig5 and Ig3 domains as compared to the vertebrates wherein, CD80-like C2-set Ig2 domains are significantly more. For example, the sticks and stones protein in *Drosophila* is composed of two-Ig5, four-Ig3, and only three-CD80-like C2-set Ig2 domains (Fig.2). Interestingly, we noticed that in both *Octopus* (Mollusca) and *Branchiostoma* (Cephalochordate) nephrin sequence is devoid of FN3 domain, though it has Ig domains (Fig.2).

**Figure 2:**
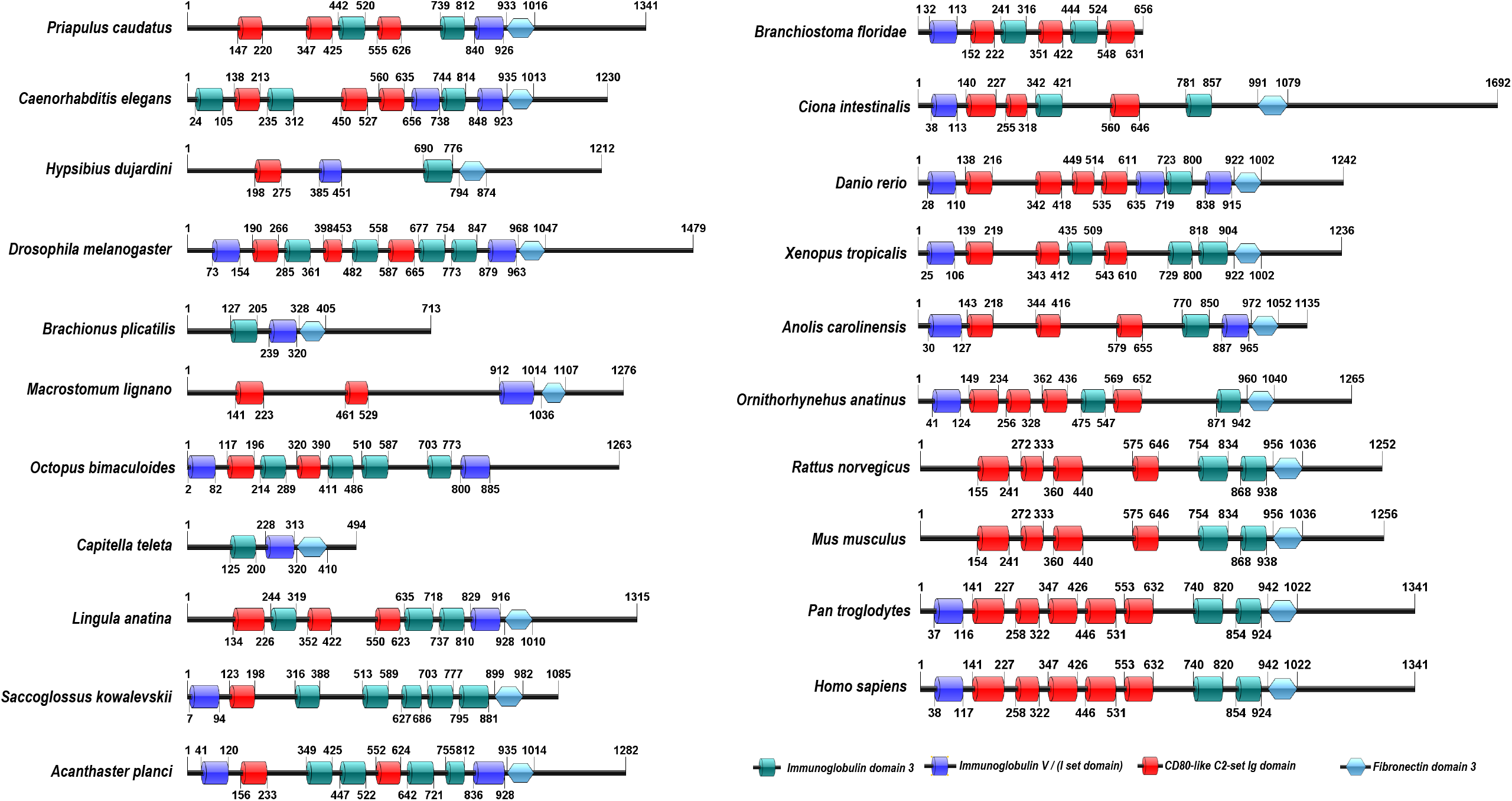
Domain organization in nephrin and its orthologs. Different colors are used to represent the subsets of Ig domains namely: Ig3 as green, CD80-like C2-set as red, and IgV as blue.

The human CD2AP protein consist of three Src Homology-3 (SH3) domains [14]. Pfam analysis of the CD2AP and the CINDR sequences showed that the three SH3 domains are highly conserved (Fig.3). Our analysis could not predict any other domains in either CD2AP or in the orthologs of CD2AP (Fig.3). Podocin in humans consist of only an SPFH domain while the rest of the sequence does not contain any known domains (Fig.4) [8]. Similar to the human podocin sequence, the podocin sequences from the other vertebrate organisms also had the SPFH domain suggesting that this domain is highly conserved among vertebrates (Fig.4). The human TRPC6 sequence consists of three conserved ankyrin repeats, a TRP domain, and an ion transport domain [15]. Our analysis of the TRPC6, TRPC3, and TRPC7 sequences showed that Ankyrin repeats, TRP, and ion transport domains were well conserved with a few exceptions (Fig.5). In the TRPC6 sequence of *Rhincodon typus* we observed only the ion transport domain but not the ankyrin repeats and TRP domain, whereas, in *Acanthaster planci* the TRPC3 protein lacked the ankyrin repeats but possessed the ion transport and the TRP domains (Fig.5).

**Figure 3:**
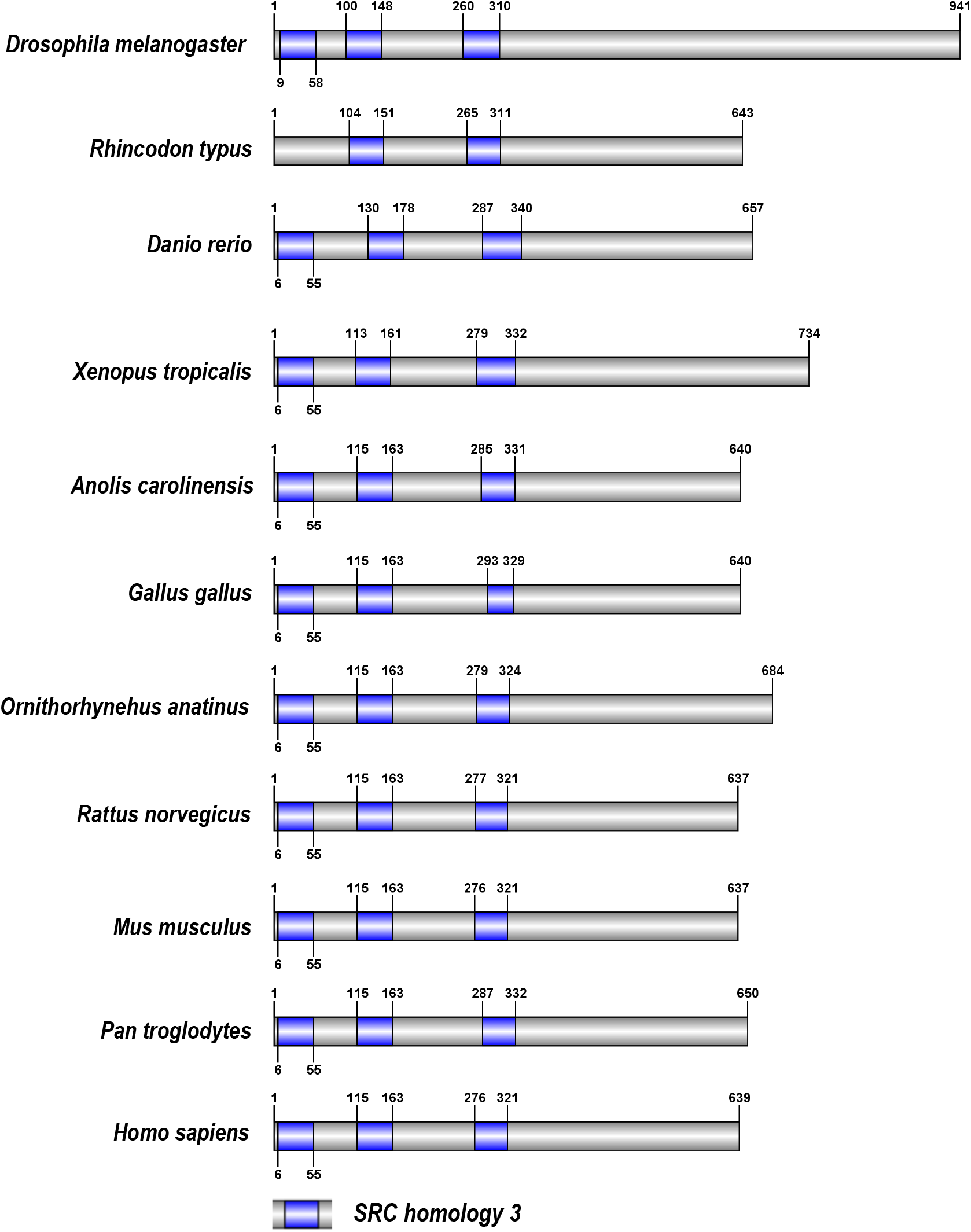
Domain organization in CD2AP and CINDR sequences.

**Figure 4:**
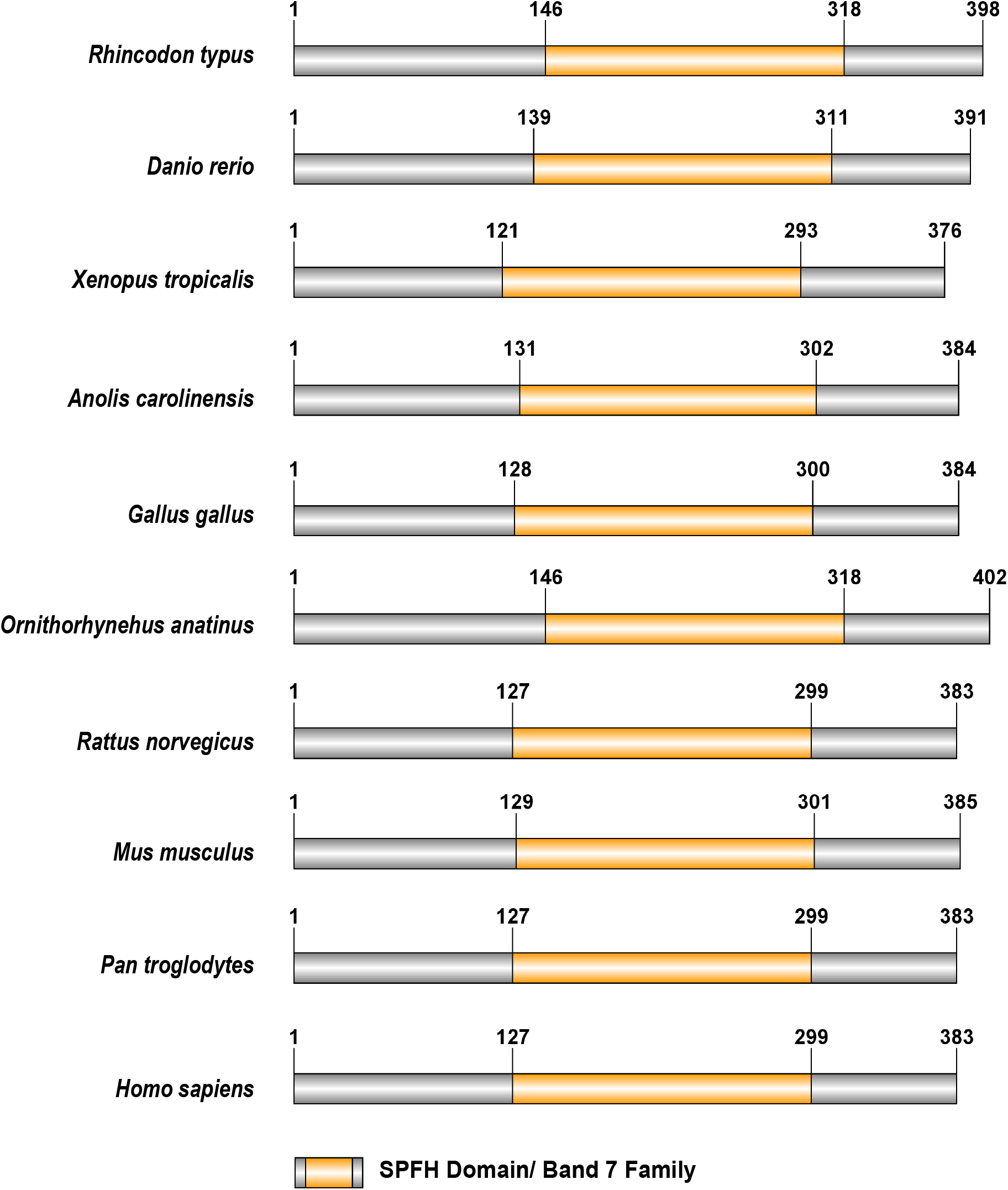
Domain organization of podocin sequences of the chordate phylum

**Figure 5:**
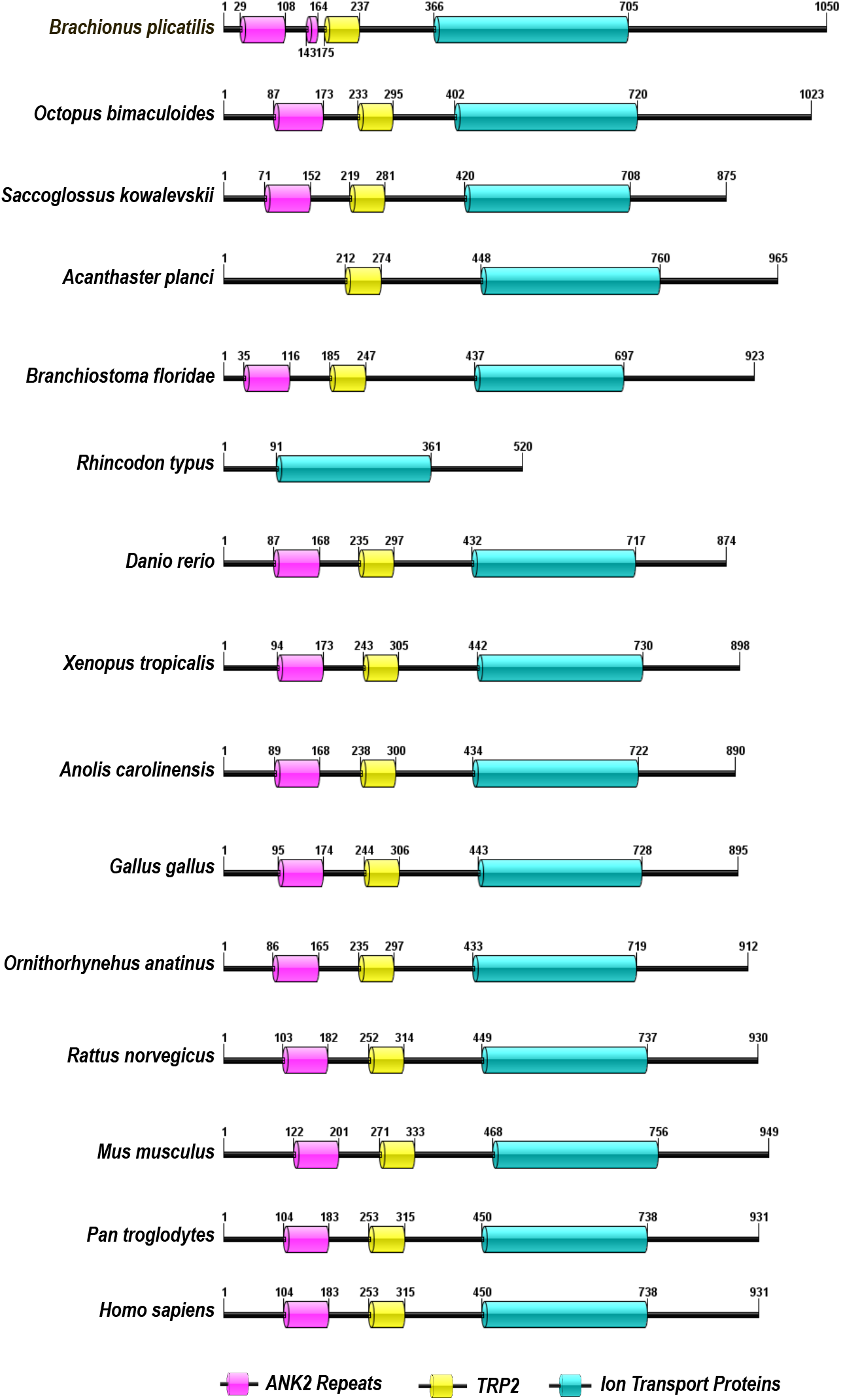
Domain organization in TRPC6, TRPC3, and TRPC7 sequences. The ankyrin repeats in the sequences are represented as pink rectangle, similarly yellow rectangle represents TRP domains, and teal represents ion transport domain.

### IURs are conserved motifs in SD proteins and its orthologs

SD proteins are known to form large complexes via homo- and heterophilic interactions which are believed to be mediated by IURs [2, 6, 15]. Since the SD proteins and its orthologs share similar domains and appreciable sequence similarity, we investigated if IUR motifs are conserved in the SD proteins and its orthologs. Our results showed that majority of the nephrin and its orthologous sequences have IURs at both the N- and C-terminus as well as in few Ig domains (Fig.6A & Table.S1). However, the nephrin orthologs in Rotifera, Mollusca, and Annelida had IURs only at the C-terminus and in the Ig domains but not at the N-terminus (Fig.6A). Also, our analysis revealed that the nephrin sequence from amphibians had IURs even in the FN3 domain. In the invertebrates, IURs were present mostly in the Ig3 and Ig5 domains and less in the CD80-like C2-set Ig2 domains whereas in the case of vertebrates the vice versa was observed. In the case of CD2AP and CINDR sequences, IURs constituted majority of the sequence (>50%) except for the regions that are a part of the first and the third SH3 domains (Fig.6B & Table.S1). But in the CD2AP of *Rhincodon typus* all the three SH3 domains were devoid of IURs but the rest of the sequence had IURs. Analysis of the podocin sequences from the Chondrichthyes to Mammalia showed IURs at both the N- and the C-terminuses (Fig.6C & Table.S1). The distribution pattern of IURs in TRPC6 and its orthologs was similar to nephrin and its orthologs. Our results indicate that TRPC6 and TRPC6 orthologs possess IURs at both the N- and C-terminus with some intermittent regions in the sequences (Fig.6D & Table.S1). Nevertheless, the TRPC6 sequences in vertebrates (Amphibia-Mammalia) the IURs were restricted only to the N- and the C-terminuses. These results suggest that IURs are conserved motifs among the SD proteins and its orthologs.

**Figure 6:**
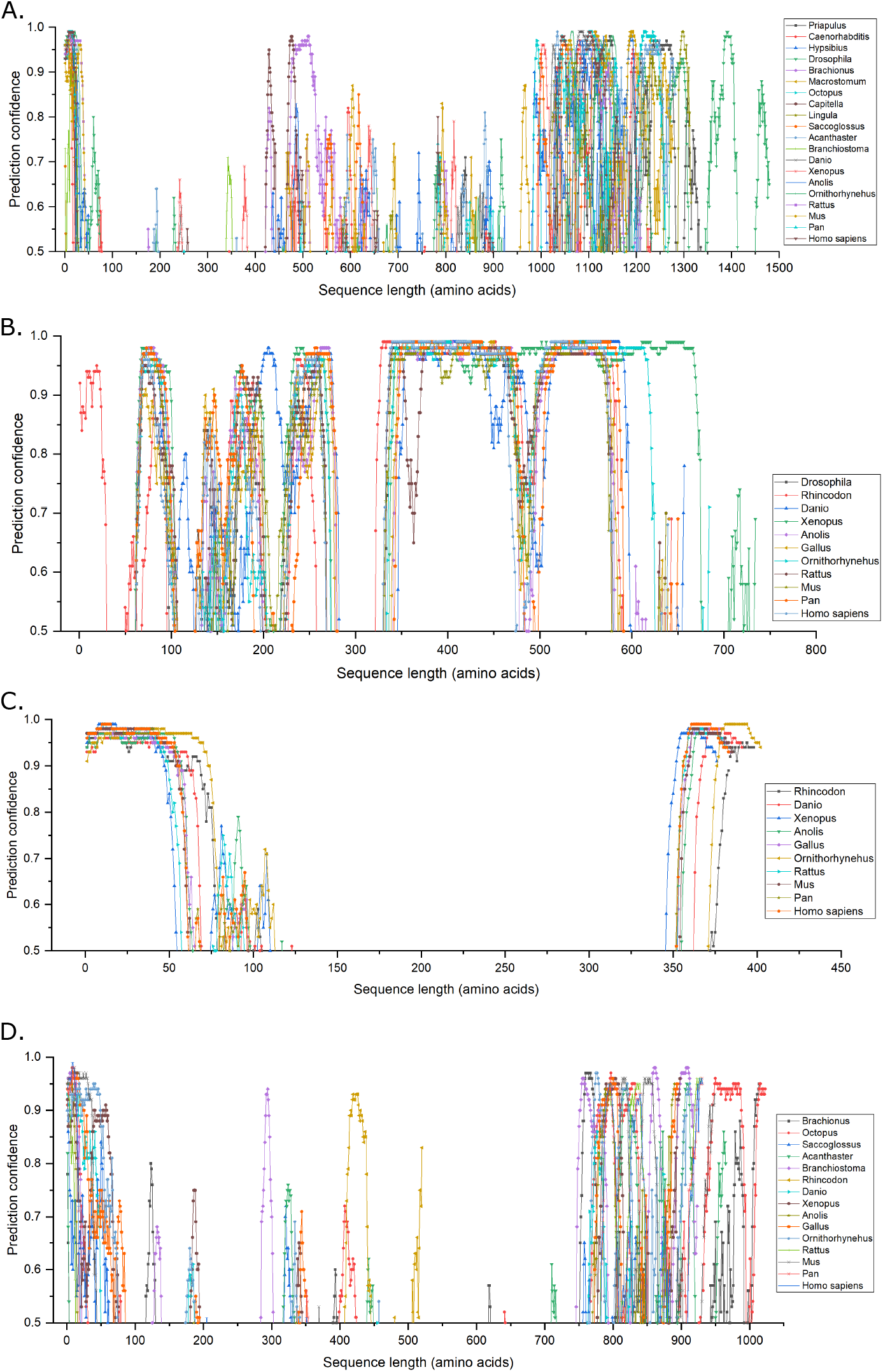
Intrinsically disordered regions (IURs) predicted by PSI-PRED for the slit-diaphragm proteins and its orthologs in various phyla: A) IURs in nephrin and orthologs, B) IURs in CD2AP and orthologs, C) IURs in podocin, D) IURs in TRPC6 and orthologs The Y-axis has been adjusted for enhanced clarity of the residues that predicted as IURs. A prediction value of ≥0.5 for a residue indicates a residue to be intrinsically unstructured.

## Discussion

Primitive nephron like structures were identified in many invertebrate phyla suggesting an evolutionary link between the vertebrate and invertebrate excretory units which further suggests that orthologous proteins similar to the vertebrate SD proteins may be present in other metazoans. Therefore, in this study, we investigated the molecular evolution of the major human podocyte SD proteins namely nephrin, CD2AP, podocin, and TRPC6 proteins that play a crucial role in the aiding the SD integrity. Our study showed that though nephrin is distributed across several phyla in both vertebrates and invertebrates, while the remaining three proteins we investigated are largely confined to vertebrates. SD proteins and its orthologs share several conserved domains and IURs indicating that the invertebrate SD proteins and its invertebrate orthologs are evolutionarily related.

In humans, nephrin is 1241 amino acids transmembrane protein made up of eight Ig domains and an FN3 domain [15]. Nephrin along with KIRREL1 forms the characteristic zipper-like structure bridging the gap between the adjacent foot process [16]. Mutations or knockdown of the gene encoding the nephrin caused Finnish-type congenital nephrotic syndrome and improper development of coronary arteries in human and mice embryos [13, 17]. Nephrin expression was also observed in the pancreatic islet cells, beta cells and lymphoid tissues [18-20]. Therefore, the divergent presence of nephrin in metazoans is observed in our analysis. Nevertheless, while analyzing the domains of the orthologs of nephrin, we noticed that in invertebrates the CD80-like C2-set Ig domains were less as compared to the nephrin sequences in vertebrates. It is known that nephrin forms dimers with nephrin/KIRREL1 from adjacent podocyte foot process [16, 21, 22]. We speculate that the increased number of CD80-like C2-set Ig domains in vertebrates may facilitate stronger homophilic interactions between neighboring nephrin/KIRREL1 molecules when compared to Ig3 or Ig5 domains. Apart from providing integrity to the SD, nephrin along with KIRREL1 also participates in signaling transduction. The cytoplasmic tails of both nephrin and KIRREL1 consist of several conserved tyrosine residues that undergo phosphorylation by Fyn (an Src family nonreceptor protein tyrosine kinase) [23]. This phosphorylation step is essential for the proper function of the SD, since deletion of the gene encoding Fyn resulted in abnormal filtration, podocyte foot process effacement, and proteinuria. Furthermore, binding of podocin with nephrin augments nephrin signaling [23, 24].

Human CD2AP is a 639 residues protein and primarily identified as an actin-binding cytoplasmic ligand for CD2 in T-cells and natural killer cells [25-27]. Localization studies showed CD2AP expression at the SD along with nephrin and podocin [9, 28]. CD2AP acts as an acting binding adaptor protein and helps in nephrin/KIRREL1 signaling in podocytes [29]. CD2AP knockout mice developed nephrotic syndrome with heavy proteinuria suggesting the importance of CD2AP in SD integrity and podocyte permselectivity [30]. Although CD2AP shares ∼50% similarity with CIN85, which belong to SH3 domain-containing kinase-binding protein 1 (SH3KBP1), our analysis identified only one ortholog for CD2AP in *Drosophila* and it is known as CINDR The adaptor molecules characteristically consist of three SH3 domains, proline rich motif, and a coiled-coil region [14]. Similarly, we observed the signature SH3 domains in CD2AP. Interestingly, CINDR also consists of SH3 domains. Similar to CD2AP, CINDR is involved in cell adhesion, cytoskeleton dynamics, and synaptic vesicle trafficking in *Drosophila* [31, 32].

Positional cloning identified *NPHS2* encodes a podocyte exclusive 383 residues integral membrane protein called podocin [8]. Podocin acts as a scaffolding molecule and provides structural integrity to the SD by forming a macromolecular complex with nephrin, CD2AP, TRPC6, and KIRREL1 [9, 33]. It was reported that besides forming a heteromeric complex, podocin like its family members associates into higher order oligomers [6, 34]. It is suggested the that the numerous interactions formed by podocin with its client proteins may be attributed to the podocin’s oligomeric nature since, each molecule in the podocin homo-oligomer can interact with one of its client proteins [15]. Podocin despite showing close homology (∼40%) with the stomatin proteins, our study could not identify the orthologs of podocin outside the vertebrate phylum. Further, the homologous region is restricted only to the SPFH domain, while the rest of the sequence did not correlate with any known domains.

The human TRPC6 is a 931 residues calcium ion transport channel protein associated with smooth muscle contraction, pulmonary endothelial permeability, neuronal protection against ischemia, and also in the structure and function of podocytes [35]. TRPC6 and its related TRPC channels are a part of a larger family of TRP proteins involved majorly in chemo- and mechanosensation [36]. It was shown that the mutations in TRPC6 caused the late onset of the focal segmental glomerulosclerosis. In some cases, the mutations increased calcium ion influx into the cell leading to altered podocyte morphology, however, the mechanistic insights of how this occurs is still poorly understood [37]. Based on the sequence similarity, the TRPC6, TRPC3, and TRPC7 proteins share appreciable homology. Furthermore, the TRPC proteins share several conserved regions namely; a) Ankyrin (ANK) repeats, b) coiled-coil domain, c) a 25 residues TRP domain, d) proline-rich sequence, followed by e) a calmodulin and IP3 receptor-binding region (CIRB region), and f) C-terminal coiled-coiled domain [37]. Therefore, it is predictable that the reciprocal best-hit method retrieved TRPC3 and TRPC7 proteins as the orthologs of TRPC6 and also that these sequences share considerable conserved domains. TRPC6 besides being a component of the slit-diaphragm, is a critical controller of the actin cytoskeleton. Activation of TRCP6 leads to the surge in cytoplasmatic Ca^2+^ levels therefore TRPC6 overexpression leads to loss of actin stress fibers in podocytes, disruption of focal adhesions, and proteinuria in mice [38, 39]. In case of nephrin, podocin, CD2AP mutations are associated with diminished function and proteinuria. Decreased expression of nephrin and podocin manifest in pathology phenotype. Alternatively, elevated TRPC6 expression is associated with glomerulopathy [39].

In vertebrates, the SD proteins are known to co-localize at the SD while, interact and form a large macromolecular complex [40]. Further, it was suggested that SD proteins consist of IURs through which these proteins may be forming complexes. IURs are areas in the protein sequences that do not adopt any secondary structure conformation in isolation, however, in the presence of an interacting partner or a suitable ligand, IURs adopt ordered structures [41]. Furthermore, they are known to play a crucial role in mediating protein-protein interactions and signaling events [42]. Since, SD proteins are predicted to have IURs, we were interested to know whether IURs are also conserved in the SD orthologs. Our results have shown that IURs are also conserved motifs across the SD orthologs which further ascertains the evolutionary relationship between SD proteins and its orthologs. Further, based on the location of IURs in nephrin, podocin, and TRPC6 it is possible that IURs may promote both homomeric and heteromeric interactions.

In conclusion, our study gives novel insights into the evolutionary relation between the vertebrate SD proteins and the invertebrate orthologs. We propose that the SD proteins may have evolved from the orthologs sequence identified in the invertebrate phyla. We also show that the unique domains present in the SD proteins are highly conserved. Further, our study shows that IURs are highly conserved motifs among the vertebrate and the invertebrate sequence which further adds evidence for the role of IURs in the SD complex formation. Though, studies reveal these four proteins interact and maintain the SD architecture, the precise stoichiometry of these proteins yet to be unraveled. Also, how mutation alters structure and functional relationship needs to be explained.

## Acknowledgements

We would like to thank Ms. Hita soni, for her feedback on our work, also SKMN would like to acknowledge Indian council of medical research for providing senior research fellowship.

## Funding Body

Indian Council of Medical research, **Grant no:** 5/4/7-9/19/NCD-II

## Conflict of interest

The authors have no conflict of interest to declare.

## Supplementary tables

**Table S1:** Amino acid residues predicted as Intrinsically unstructured regions (IURs) and IUR- binding domain (BD) by the PSI-PRED server in the slit-diaphragm proteins and its orthologs in various metazoans.

## Notes

### Competing Interest Statement

The authors have declared no competing interest.

