## Supplementary material for "Molecular Evolution of Podocyte Slit-diaphragm Proteins": Table.S1

**Table S1:** Amino acid residues predicted as Intrinsically unstructured regions (IURs) and IUR-binding domain (BD) by the PSI-PRED server in the slit-diaphragm proteins and its orthologs in various metazoans.

| **Intrinsically unstructured regions (IURs) predicted in nephrin and its orthologs** | | | |
| --- | --- | --- | --- |
| **Names of the organisms** | **NCBI**  **Accession ID** | **IURs** | **IUR-(Binding domains)** |
| *Priapulus caudatus* | XP_014667088.1 | 833;835-839;882-883;889;1030-1036;1078-1157;1197-1283;1300-1314;1316;1318-1322;1325-1326;1330-1331 | 01-24;834;840-844;  1029;1037-1044;1315;1317;1323-1324;1327-1328;1335 |
| *Caenorhabditis elegans* | NP_001309674.1 | 74-75;345;482-492;544-551;589-593;595-600;754-756;843;886-887;891-892;1026-1028;1082-1126;1224 | 01-17;71-73;76-78;594;601-604;883-885;889-890;1025;1029-  1040;1073-1081;1184-1201;1125;1228-1230 |
| *Hypsibius dujardini* | [OQV25698.1](https://www.ncbi.nlm.nih.gov/protein/OQV25698.1?report=genbank&log$=prottop&blast_rank=1&RID=VJ5MSDJ9014) | 06-30;32-33;36;42-46;459;461;623-629;696-699;702-704;739;741-746;880-882;889-891;894;981-988;990-999;1027-1200 | 01-05;31;34-35;37-41;48-50;451-456;630-  641;883-888;892-893;895;989;1000-1012 |
| *Euperipatoides rowelli* | No relevance | | |
| *Drosophila melanogaster* | NP_788286.1 | 775-781;912-922;1053-1074;1111-1112;1130;1132-1133;1138-1139;1141-1167;1178-1180;1187;1189-1191;1208-1213;1226;1266-1310;1345;1349-1412;1452-1456;1461-1472 | 01-25;47-51;53-73;228-232;782-787;923;1075;1110;1113-1129;1137;1140;1173-1176;1188;1214-1225;1227-1232;1450-1451;1457-1460;1473-1479 |
| [*Brachionus plicatilis*](https://blast.ncbi.nlm.nih.gov/Blast.cgi#alnHdr_RNA22476) | [RNA22476.1](https://www.ncbi.nlm.nih.gov/protein/RNA22476.1?report=genbank&log$=prottop&blast_rank=1&RID=VU6MM62J014) | 175-176;421-423;470-573 | 424-434;576-580;618-624;630-633 |
| *Macrostomum lignano* | [PAA62947.1](https://www.ncbi.nlm.nih.gov/protein/PAA62947.1?report=genbank&log$=prottop&blast_rank=1&RID=VJ6A3YY9014) | 24;477-481;554;557;559;596-612;614-616;618;628-630;687-694;697;785-797;846-847;849-857;862-863;956-975;1118-1119;1121;1145-1150;1178-1276 | 01-22;463-474;555-556;558;560-563;695-696;798-800;858-861;864;1123-1144;1151 |
| *Octopus binaculoides* | XP_014776688.1 | 551-795-797;986-987;1034-1106;1111-1112;1116;1126-1127;1129;1195-1260 | 983-985;988-1000;1132-1141;1261 |
| *Capitella teleta* | ELU18150.1 | 426;449 | 421-425;427-442;466-494 |
| *Notospermus geniculatus* | No relevance | | |
| *Lingula anatina* | [XP_013380213.1](https://www.ncbi.nlm.nih.gov/sites/entrez?cmd=Search&db=protein&term=XP_013380213.1&dopt=GenBank) | 580;582;585;587-588;681-688;877;1020-1024;1073-1104;1124-1126;1151-1153;1164;1222;1229;1232-1236;1239-1257;1287-1288;1298;1301-1309 | 01-24;581;583-584;586;589-590;1025-1033;1066-1072;1128-1133;1142-1149;1163;1165-1166;1223-1228;1230-1231;1237-1238;1289-1297;1299-1300;1310-1315 |
| *Phoronis australis* | No relevance | | |
| *Saccoglossus kowalevskii* | XP_006819647.1 | 01-02;548-561;611-616;843;992-1000;1008 | 562;617-626;1001-1007;1009-1020;1039-1062;1083-1085 |
| *Acanthaster planci* | XP_022110423.1 | 186-188;192-197;360-361;586-594;645-653;657;659-690;695;749;751;832-839;876-878;1029-1030;1078-1183;1197;1204-1281 | 01-31;190-191;595-  596;654-655;840-841;879-889;890;1026-  1028;1031-1042;1186-1188;1198-1201;1282 |
| *Branchiostoma floridae* | XP_002590121.1 | 04-29;350-351;653-656 | 01;339-349;386 |
| [*Ciona intestinalis*](https://blast.ncbi.nlm.nih.gov/Blast.cgi#alnHdr_XP_002122747) | [XP_002122747.1](https://www.ncbi.nlm.nih.gov/protein/XP_002122747.1?report=genbank&log$=prottop&blast_rank=1&RID=VU4SEMTZ014) | Sequence too long | |
| *Petromyzon marinus* | No relevance | | |
| *Rhincodon typus* | No relevance | | |
| *Danio rerio* | XP_017206503.1 | 240-244;581-584;823-828;870-871;874;1017-1018;1076-1129;1185-1230 | 01-22;245-247;829-  832;872-873;875-880;1019-1035;1070-1075;1132-1135;1140 |
| *Xenopus tropicalis* | XP_017944912.1 | 73;75-76;236;239-242;373-375;382;475-480;483-485;575-576;580-581;633-637;811-818;823;1000-1001;1003-1005;1018-1019;1021-1023;1029;1075;1079-1080;1082-1083;1085 | 01-20;243-246;376- 381;383;638-645;819-821;994-999;1002;  1006-1017;1020 |
| *Anolis carolinensis* | XP_016851514.1 | 478-482;488;491-493;621-624;627;811-817;820;922-923;1059  1060;1064-1100;1102-1107 | 01-23;483-487;489-490;617-620;1061;1101;1108-1113;1115-  1135 |
| *Gallus gallus* | Sequence not available | | |
| *Ornithorhynehus anatinus* | XP_028921034.1 | 06-08;191-193;196;457-459;614-615;617-618;674-677;679;861-863;867-870;909-912;916-917;1108-1109;1111-1165 | 01-05;09-35;1055-1075;1110 |
| *Rattus norvegicus* | NP_072150.1 | 06-09;39;41-42;502-514;607;795;800-804;1050-1052;1054;1075-1076;1115-1117;1120-1150;1197 | 01-05;10-38;40;796-799;1053;1055-1071;1103-1114;1118-1119;1152-1156;1159-1173;1206-1209 |
| *Mus musculus* | NP_062332.2 | 06-09;13-15;17;501-515;604;606-608;610-611;669-670;795-796;799-804;1052;1070-1071;1112-1157 | 01-05;10-12;16;18-41;797-798;1050-1051;1053-1069;1104-1111;1159-1178 |
| *Pan troglodites* | XP_016791216.2 | 250;495-500;590-593;650;786-791;843-847;891-894;898-900;1029-1030;1034-1039;1042-1044;1052-1059;1099-1136 | 01-25;781-785;848-851;1040-1041;1045-1051;1091-1098;1139-1143;1146-1164 |
| *Homo sapiens* | NP_004637.1 | 257-258;493-502;589-593;596-597;649-655;787-790;891-893;898-899;1028-1031;1033-1044;1055-1057;1101-11281131-1137 | 01-27;656-657;780-786;1032;1045-1054;1090-1100;1129-1130;1141-1143;1146-1165 |
| **Intrinsically unstructured regions (IURs) predicted in CD2AP and its orthologs** | | | |
| *Priapulus caudatus* | No relevance | | |
| *Caenorhabditis elegans* | No relevance | | |
| *Hypsibius dujardini* | No relevance | | |
| *Euperipatoides rowelli* | No relevance | | |
| *Drosophila melanogaster* | XP_016866130.1 | 62-68;85-97;127-132;160-198;219-267;333-476 | 69-84;133-154 |
| *Brachionus plicatilis* | No relevance | | |
| *Macrostomum lignano* | No relevance | | |
| *Octopus bimaculoides* | No relevance | | |
| *Capitella teleta* | No relevance | | |
| *Notospermus geniculatus* | No relevance | | |
| *Lingula anatina* | No relevance | | |
| *Phoronis australis* | No relevance | | |
| *Saccoglossus kowalevskii* | No relevance | | |
| *Acanthaster planci* | No relevance | | |
| *Branchiostoma floridae* | No relevance | | |
| *Ciona intestinalis* | No relevance | | |
| *Petromyzon marinus* | No relevance | | |
| *Rhincodon typus* | XP_020370685.1 | 13-14;51-52;58;157-205;227-234;239-257;322-587;589-590;636 | 01-12;15-29;50;54-57;59-62;68-95;235-238;642-643 |
| *Danio rerio* | NP_001008583.2 | 62-124;139-142;159-169;172-282;346-599;649-650 | 143-158;655-657 |
| *Xenopus tropicalis* | NP_001121435.1 | 61-105;155-204;206-208;211-212;218-273;337-675;705;712;726-727 | 141-153;708-711;713-724;733-734 |
| *Anolis carolinensis* | XP_008114796.1 | 62-105;145-199;224-278;341-477;483-484;486;488-581;604-607;611;615;632-633 | 133-143;639-640 |
| *Gallus gallus* | NP_001305332.1 | 62-106;128-203;222-277;339-584;586-587;632-633 | 639-640 |
| *Ornithorhynehus anatinus* | XP_028928027.1 | 62-103;133-135;137-139;146-147;151-203;224-272;334-624;677 | 140-142;148;683-  684 |
| *Rattus norvegicus* | AAM47029.1 | 62-106;137-138;166-201;203;224-268;331-578;629;630 | 140-149;636-637 |
| *Mus musculus* | AAI38375.1 | 62-104;144;161-210;213;215-267;331-578;629-630 | 146-151;636-637 |
| *Pan troglodytes* | XP_009449690.1 | 62-104;126-190;210-211;231-280;342-483;493-494;499-591;642-643 | 649-650 |
| *Homo sapiens* | NP_036252.1 | 62-102;147;149;170-198;223-268;331-478;481-  580;631-632 | 127-142;148;638-640 |
| **Intrinsically unstructured regions (IURs) predicted in podocin and its orthologs** | | | |
| *Priapulus caudatus* | No relevance | | |
| *Caenorhabditis elegans* | No relevance | | |
| *Hypsibius dujardini* | No relevance | | |
| *Euperipatoides rowelli* | No relevance | | |
| *Drosophila melanogaster* | No relevance | | |
| *Brachionus plicatilis* | No relevance | | |
| *Macrostomum lignano* | No relevance | | |
| *Octopus bimaculoides* | No relevance | | |
| *Capitella teleta* | No relevance | | |
| *Notospermus geniculatus* | No relevance | | |
| *Lingula anatina* | No relevance | | |
| *Phoronis australis* | No relevance | | |
| *Saccoglossus kowalevskii* | No relevance | | |
| *Acanthaster planci* | No relevance | | |
| *Branchiostoma floridae* | No relevance | | |
| *Ciona intestinalis* | No relevance | | |
| *Petromyzon marinus* | No relevance | | |
| *Rhincodon typus* | XP_020382509.1 | 02;05-78;105 | 01;03-04;100-104;372-398 |
| *Danio rerio* | NP_001018155.2 | 3-18;21;24-69;101;104-105;123 | 1-2;19-20;22-23;93-98;363-391 |
| *Xenopus tropicalis* | XP_017949096.1 | 09-17;31-34;36-54;348 | 01-08;18-30;35;75-87;103-110;346-347;349-376 |
| *Anolis carolinensis* | XP_003225506.1 | 06-11;13;17-26;30-39;46-64;117 | 01-05;12;14-16;27-29;40-45;83-98;355-384 |
| *Gallus gallus* | XP_422265.3 | 08-11;13;18-19;21-25;34-37;39-65;83;95;353-358 | 01-07;12;14-17;20;26-33;38;88-94;96;359-384 |
| *Ornithorhynehus anatinus* | XP_001515734.2 | 01-79;81-82;84;383-384 | 85-86;98-112;371-382;385-402 |
| *Rattus norvegicus* | NP_570841.2 | 18;29-32;34-57;92-94 | 01-17;19-28;33;76-90;353-383 |
| *Mus musculus* | NP_569723.1 | 07-08;16-23;28-62;97 | 01-06;09-15;24-27;354-385 |
| *Pan troglodytes* | XP_016788875.1 | 06-13;68;80-83;18-24;  26-61;94 | 01-05;14-17;66-67;86-93;95-96;352-383;25 |
| *Homo sapiens* | NP_055440.1 | 06-13;16;18-24;26-61;  67-68;94;352;364-366 | 01-05;14-15;17;25;66;80-83;86-93;95-96;353-363;367-383 |
| **Intrinsically unstructured regions (IURs) predicted in TRPC6 and its orthologs** | | | |
| Priapulus caudatus | No relevance | | |
| *Caenorhabditis elegans* | No relevance | | |
| *Hypsibius dujardini* | No relevance | | |
| *Euperipatoides rowelli* | No relevance | | |
| *Drosophila melanogaster* | No relevance | | |
| *Brachionus plicatilis* | RNA42410.1 | 01-02,117,122-129,390-396,618-620,754-777,803,805-807,810,943,963,997-998 | 116,118-121,752-753,808-809,945-950,956-961,967-992,1003-1015 |
| *Macrostomum lignano* |  | No relevance | |
| *Octopus bimaculoides* | XP_014783218.1 | 07-08,12,15-19,23-26,28-29,31-76,78-79,399-424,641-642,766-840,866,870,926,928,932-994,1000-1001 | 01-06,09-11,13-14,20-22,27,30,897-907,1004-1023 |
| *Capitella teleta* | No relevance | | |
| *Notospermus geniculatus* | No relevance | | |
| *Lingula anatina* | No relevance | | |
| *Phoronis australis* | No relevance | | |
| *Saccoglossus kowalevskii* | XP_002730569.1 | 06-07,18-41,319-328,  764-765,799,846 | 01-05,08-17,43-61,756-759,830-843,869-875 |
| *Acanthaster planci* | XP_022103606.1 | 02-03,317-333,440-448,810-841,869,871-873,  879-880,906 | 01,710-716,807-809,863-868,870,874-877,893-  905,907-920,950-965 |
| *Branchiostoma floridae* | XP_002607434.1 | 125,133,135-138,284-  302,746-793,820-821,  824-826,828,887-888,  890,898-906,909,914-  916 | 01-21,128-132,134,827,851-874,889,891-897,907-908,910-913,917-923 |
| *Ciona intestinalis* | No relevance | | |
| *Petromyzon marinus* | No relevance | | |
| *Rhincodon typus* | XP_020383938.1 | 410-441,443-445,477 | 409,442,509-520 |
| *Danio rerio* | XP_005161247.1 | 06-49,178-186,455,778-788,790-791,820-821,823-825,867 | 01-05,763-777,826-828,872-874 |
| *Xenopus tropicalis* | XP_002935616.2 | 01-68,73-75,181-194,  337-344,777-807,811-  816,843-844,846-848,  851 | 71-72,808-810,849-850,881-898 |
| *Anolis carolinensis* | XP_008106273.1 | 07-08,186,336,773-793,832,834-840,845-846,  848 | 01-06,09-15,770-772,  794-806,841-844,872-  890 |
| *Gallus gallus* | XP_417184.4 | 01-67,188-194,339,341-351,780-813,841-842,  844-847,849 | 70-85,773-779,848,874-895 |
| *Ornithorhynehus anatinus* | XP_028903565.1 | 02-74,174-187,330,333-337,341,457,772-773,  777-788,827,829,854,  859,867 | 01,764-771,774-776,  801-826,828,856-858,  860-865,893-912 |
| *Rattus norvegicus* | NP_446011.1 | 350,789-792,795-800,  802-807,812,847,874-  879,881-883 | 01-12,782-788,793-794,801,813-816,818-846,  910-930 |
| *Mus musculus* | XP_006509912.1 | 06-13,15-26,28-29,31-  43,369,808-810,827,830-836,838-839,848-866,905 | 01-05,14,27,30,804-807,811-826,837,840-847,  893-902,930-949 |
| *Pan troglodytes* | XP_016777341.2 | 351-353,804-807,814-  816,818,827-837,845-  846,848 | 01-17,783-803,819-826,  838-844,847,875-884,  910-931 |
| *Homo sapiens* | NP_004612.2 | 17,205,351-352,804-808,812-818,820-848,876-880,882,884 | 01-16,18-19,783-803,819,875,883,910-931 |
